# Statistical Inconsistency of Error-correction Objectives for Perfect Phylogenies

**DOI:** 10.64898/2026.07.20.739615

**Authors:** Gryte Satas, Matthew A. Myers, Sohrab P. Shah

## Abstract

The binary perfect phylogeny, in which each mutation arises exactly once on an evolutionary tree and is never lost, is a well-studied idealized phylogenetic model. When observed data has errors, a common approach to phylogeny inference is to seek a tree that minimizes the number of error corrections (“flips”) needed to fit a perfect phylogeny. These objectives draw on an intuitive justification: minimizing implied errors should prefer the true tree in expectation.

We test this assumption using a generative model with independent errors and prove that error-correction objectives are statistically inconsistent for all positive error rates: in expectation, the minimum-cost tree need not be the true tree. Our proof is constructive and yields counterexamples involving any tree topology and any positive error rates, demonstrating the ubiquity of the phenomenon. The core problem is that tree topologies can explain observed mutation patterns with fewer errors than actually occurred, and differ in their ability to do so, introducing systematic bias. This mechanism is distinct from previously identified sources of inconsistency such as homoplasy or incomplete lineage sorting, since the error-free setting is trivially consistent for perfect phylogenies.

We investigate how often this failure may occur in practice. Simulations calibrated to error rates from single-cell sequencing data show that an incorrect tree is preferred over the true tree in a substantial fraction of cases (over 50% in some settings) with rates increasing with tree size. Moreover, winning trees are not random but share specific topological features. Notably, at error rates typical of single-cell sequencing data, trees with deeper, more imbalanced topologies are consistently favored over more balanced ones. These results demonstrate that inconsistency is not a theoretical edge case, and that understanding when and how it arises is important when interpreting results in practice.

**Supplementary Material:** *Source code*: https://github.com/shahcompbio/phylo_inconsistency

**Funding:** This research was supported in part by NCI SPORE (P50 CA247749-01)

## 1 Introduction

The binary perfect phylogeny is a well studied model in phylogenetics in which each mutation arises exactly once on an evolutionary tree and is inherited by all descendants, with no losses or reversions [27]. This has also been referred to as the infinite-sites model, as it can be viewed as a limit of finite-sites models where the number of sites grows faster than the number of mutations, so that homoplasy and reversion become vanishingly unlikely [31]. Under this idealized model, each mutation imposes a hard compatibility constraint on the tree topology. As a result, the underlying tree topology is uniquely determined once sufficiently many informative mutations are observed, and can be found in linear time [1, 23]. In many applications, however, mutations are not observed directly but are measured through a noisy process. In single-cell sequencing data, for example, allelic dropout and limited coverage produce false negatives, while sample preparation, sequencing, and alignment artifacts introduce false positives [47]. In the presence of such errors, mutations no longer impose exact compatibility constraints, and the guarantees of the error-free perfect-phylogeny model no longer apply. This has led to a family of combinatorial formulations for phylogeny inference under measurement error [9]. These approaches seek an optimal tree that minimizes the number or weighted cost of entry-wise corrections (“flips”) to make the observed data compatible with a perfect phylogeny. The simplest case where all flips have equal cost is well known as the minimum-flip perfect phylogeny problem or the minimum-flip consensus tree problem [9]. Prior work has focused on studying the computational complexity and behavior of the problems and developing algorithms that find or approximate minimum-cost solutions [2, 5, 6, 8, 9, 10]. Applied variants of these objectives have been widely used in cancer phylogenetics, for reconstructing tumor evolutionary histories from noisy single-cell mutation data (see Related Work in Section 1.1).

Error-correction approaches are implicitly motivated by an intuition that minimizing the number of implied observation errors should recover the true tree without bias, at least when noise itself is unbiased and sufficient data are observed. However, to our knowledge, this assumption has not been systematically examined; it remains unclear whether these objectives yield statistically consistent estimators of the true tree under reasonable generative models of evolution and observation noise. Statistical consistency is a property of an estimator that guarantees convergence to the true parameter (in this case, the true tree) as the amount of data increases [21, 46]. It has been a topic of longstanding interest in phylogenetics [18, 24, 43] (see Related Work in Section 1.1); however, perfect phylogeny inference has not been examined in this context, likely because consistency is trivial in the error-free setting. The perfect phylogeny model is in this sense a natural minimal test case: it strips away all evolutionary sources of ambiguity that are known to cause phylogenetic inconsistency (e.g., homoplasy, incomplete lineage sorting), leaving only the interaction between observation noise and the objective to explain any inconsistency that arises. While few applied methods use the basic perfect phylogeny error-correction objective in its pure form, most retain the same core structure of hard-assigning mutations to branches and selecting the tree that minimizes the resulting cost. Thus, any inconsistency that arises here will likely carry through to those settings as well.

In this work, we study error-correction objectives for perfect phylogeny reconstruction under a generative model with independent observation errors. We show that these objectives are not statistically consistent for any positive error rates, even when the error penalties are perfectly calibrated to the true error rates. That is, even with known error rates and unbiased errors, the minimum-flip objective need not favor the true tree in expectation. Moreover, the objectives can be positively misleading, converging to a specific incorrect tree with increasing support as the number of mutations increases. The source of this inconsistency is not algorithmic failure, limited data, or ambiguity in the evolutionary model, but a structural interaction between observation noise and phylogenetic topology under the objective. The objective seeks a tree that explains the observed data with the fewest, or lowest cost, corrections. Consequently, errors are attributed to spurious evolutionary structure whenever doing so reduces the total cost of flips required to achieve compatibility.

Crucially, this process is not uniform with respect to tree topology: trees vary in their ability to absorb different types of errors. Certain trees can systematically explain false positive or false negative errors with fewer corrections than the true tree, leading to lower objective values in expectation. We prove that inconsistency is ubiquitous in tree space: there exist counterexamples that violate consistency for at least any pair of trees that differ by one internal edge and for any positive error rates.

Which topology is systematically favored is determined by error rates, tree topology and branch lengths; a topology may have a systematic advantage over the true tree under one branch length distribution but not another. While the theoretical proofs rely on arbitrary branch length distributions, we also demonstrate that constraining edge lengths to be equal does not eliminate inconsistency. We next performed computational analyses to investigate how often inconsistency arises under biologically motivated edge length distributions and error rates typical of single-cell sequencing data. We find that inconsistency arises in a non-trivial fraction of tested cases, reaching over 50% in some settings, with rates appearing to increase with tree size. At these error rates, the bias has a consistent directional signature toward more caterpillar-like trees. These results demonstrate that inconsistency in perfect phylogeny error-correction inference can arise in practice under non-pathological conditions.

### 1.1 Related work

#### Consistency of maximum parsimony in classical phylogenetics

Maximum parsimony is one of the oldest and most widely used approaches to phylogenetic reconstruction [22], seeking the tree that minimizes the total number of changes implied along its branches: a parsimony criterion applied directly to evolutionary events. Perfect phylogeny error-correction objectives are related in spirit but the parsimony criterion is applied to the observation error process rather than the evolutionary process.

Consistency of parsimony-based phylogenetic inference has been extensively studied [18, 24, 26, 41]. Maximum parsimony is known to be statistically inconsistent in certain parameter regimes, converging to an incorrect topology with increasing confidence as data accumulate. The phenomenon was first demonstrated by Felsenstein [18], who showed that under certain branch length configurations, long-branch attraction causes maximum parsimony to favor incorrect topologies, producing the “Felsenstein zone.” Subsequent work generalized inconsistency results to arbitrary numbers of taxa and *k*-state character models as well as demonstrated constrained regions of parameter space where parsimony is consistent [26, 41, 42, 44]. A second and distinct source of inconsistency arises under the multispecies coalescent, where various concatenation-based methods can converge to an incorrect species tree, because incomplete lineage sorting can create systematic discordance across loci [15, 14, 28, 29, 34, 36, 35, 45, 49].

In both settings, inconsistency is a consequence of the evolutionary process itself: homoplasy or lineage sorting produce systematic biases that the parsimony objective cannot overcome. The results we present here demonstrate a distinct mode of estimator failure: the perfect phylogeny assumption precludes homoplasy by construction, and a lack of recombination means that lineage sorting is not a factor.

#### Applications in cancer phylogenetics

The rooted perfect phylogeny model has been widely adopted in cancer genomics, as tumor evolution operates over a relatively short time frame that limits the accumulation of homoplasy, and the normal genome provides a known ancestral state [4]. With the rise of single-cell sequencing technologies, perfect phylogeny error-correction objectives have become a natural fit for inference from noisy single-cell mutation data. A range of methods have been developed in this setting, including both combinatorial and probabilistic formulations [3, 7, 16, 25, 30, 37, 38, 48, 51] as well as extensions that relax the perfect phylogeny assumption (e.g., Dollo or finite sites models), incorporate additional data modalities such as copy number, or combine multiple types of data (e.g., combining bulk and single cell) [11, 17, 20, 32, 33, 39, 40, 50, 52].

## 2 Background

### 2.1 Perfect phylogenies

A *phylogenetic tree* is a rooted binary tree *T* on *n* leaves with edge set *E*(*T*). The root *r* has a single child connected by the trunk edge *e*_*r*_; all other internal nodes have exactly two children. Each edge *e* has an associated length *p*_*e*_ > 0, normalized so that ∑_*e*∈*E*(*T*)_ *p*_*e*_ = 1.

#### ▶ Remark 1.

**Trunk edge convention**. Representing trees with a trunk edge is common and convenient in cancer phylogenetics for modeling mutations shared by all cells. Such trees can be reconciled with the standard definition of a full rooted binary tree with the addition of a dummy outgroup attached to the root that is treated as a noiseless ancestral reference.

Mutation data are represented as a binary matrix *X* ∈ {0, 1}^*n*×*m*^, where rows correspond to leaves and columns to mutations. We assume that *X* is generated under the binary *perfect phylogeny model* [23], where each mutation arises exactly once on the tree and is inherited by all descendants. Let *T* ^∗^ represent the true tree from which *X* was generated. Each mutation independently selects an edge *e* ∈ *E*(*T* ^∗^) with probability *p*_*e*_. This defines the true assignment function Φ^∗^ : {1, … , *m*} → *E*(*T* ^∗^), where Φ^∗^(*j*) is the edge on which mutation *j* arises. More generally, for any candidate tree *T* , an *assignment function* Φ : {1, … , *m*} → *E*(*T*) maps each mutation to an edge of *T* . For each edge *e* ∈ *E*(*T*), let *D*_*e*_ denote its *descendant set*, i.e. the set of leaves whose root-to-leaf path contain *e*. Under the perfect phylogeny model, an assignment Φ induces a binary mutation matrix *X*^Φ^ ∈ {0, 1}^*n×m*^ defined by 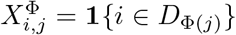, where **1**{…} denotes the indicator function. That is, all descendant leaves of *e* have state 1 for mutation *j*, and all other leaves have state 0.

#### ▶ Definition 2.

**Compatibility**. *A mutation matrix X is* ***compatible*** *with a tree T if there exists an assignment* Φ *such that X* = *X*^Φ^.

Thus, under a perfect phylogeny model with strictly positive edge lengths, each observed mutation places a hard constraint on the tree inference. Provided that sufficiently many mutations are observed to place at least one mutation on every internal edge, the true tree *T* ^∗^ is the unique tree compatible with the data [23].

#### Observation model

Given a true mutation matrix *X* ∈ {0, 1} ^*n×m*^, generated under a perfect phylogeny model, we assume that the observed matrix 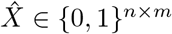 is generated under an error model where each entry of *X* is flipped independently with fixed probability. That is,

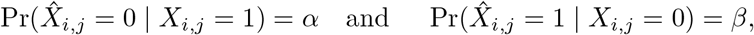

where *α* ∈ (0, 0.5) is the false negative rate and *β* ∈ (0, 0.5) is the false positive rate.

### 2.2 Minimum flip-cost perfect phylogeny problem (MFCP)

Under the observation model described above, the guarantees of the noiseless setting no longer hold: the observed matrix 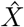 is not in general compatible with the true tree, and compatibility alone cannot guide inference. This has motivated a family of well-studied combinatorial problems for phylogeny inference for data with errors. Rather than requiring exact compatibility, these problems seek to minimize the number or cost of entry-wise corrections (“flips”) needed to make the observed data compatible with a perfect phylogeny.

For a candidate tree *T* and an assignment function Φ : {1, … , *m*} → *E*(*T*), discrepancies between the observed matrix 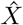 and the induced matrix *X*^Φ^ take two forms: 0 → 1 flips (entries where 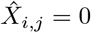 but 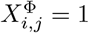) and 1 → 0 flips (entries where 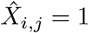 but 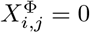). Let *w* , *w >* 0 denote the costs of 0 → 1 and 1 → 0 flips, respectively. Note that the flip directions are defined relative to editing the observed matrix 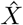 into a matrix compatible with a candidate tree, and should not be confused with the directions of the underlying observation errors. Likewise, *w*_*P*_ and *w*_*N*_ are user-defined parameters of the inference objective, and should not be confused with the latent error rates *α* and *β* in the generative model. While *w*_*P*_ and *w*_*N*_ can be set to reflect the relative costs of false positives and false negatives, we make no assumptions about their relationship in the theoretical analyses that follow.

For two binary vectors ***x***, 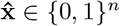, the flip cost is

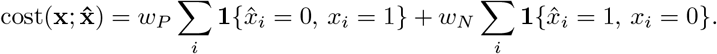

For any edge *e* ∈ *E*(*T*), let **x**_*e*_ ∈ {0, 1}^*n*^ denote the mutation pattern induced by *e*, where the 1-set of **x**_*e*_ corresponds to the descendant set *D*_*e*_. The *flip cost* of a tree *T* with respect to an observed matrix 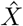 is defined as

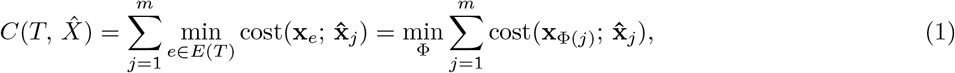

where 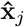 is the *j*-th column of 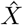, and the minimum is taken over all assignment functions Φ : {1, … , *m*} → *E*(*T*). Note that 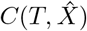 is invariant under multiplying both *w*_*P*_ and *w*_*N*_ by the same positive constant, and as such there is only one degree of freedom in the choice of weights, which can be parameterized by the ratio *w*_*P*_ */w*_*N*_ . We will continue to refer to them as two separate parameters, as it provides a more intuitive interpretation. Throughout, *w*_*P*_ and *w*_*N*_ are treated as fixed parameters of the objective and suppressed from the notation of *C* and all quantities derived from it.

This yields the following computational problem:

#### ▶ Problem 3.

***Minimum flip-cost perfect phylogeny problem (MFCP)***. *Given an observed binary matrix* 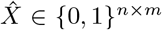 *and weights w*_*P*_ , *w*_*N*_ > 0, *find a tree T minimizing the flip cost* 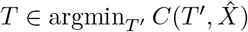.

Specific choices for weights *w*_*P*_ and *w*_*N*_ yield well-studied special cases: the minimum flip problem (MFP, *w*_*P*_ = *w*_*N*_ = 1), the insertion-only flip problem (IFP, *w*_*N*_ ≫ *w*_*P*_ , so 1 → 0 flips are never used), and the deletion-only flip problem (DFP, *w*_*P*_ ≫ *w*_*N*_ , so 0 → 1 flips are never used) [9].

▶ Remark 4. **Relationship to maximum likelihood inference**. A natural response to inconsistency of a combinatorial objective is to switch to a maximum likelihood formulation. However, jointly maximizing likelihood over trees and mutation assignments (i.e., 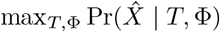) under the observation model described above yields an objective that falls within the MFCP family, so the inconsistency results here apply directly. Specifically, when error rates are assumed to be uniform and independent, the log-likelihood decomposes into a sum of per-entry penalties, and maximizing it is equivalent to minimizing a weighted flip cost with weights 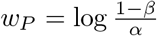 and 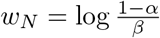 (see Appendix A.4 for details). Some approaches use entry-specific likelihoods incorporating additional information such as read counts; these can be viewed as generalizations of the MFCP objective in which the i.i.d. case is a special case. As they still rely on hard-assigning mutations to edges and optimizing a sum of penalties, the inconsistency mechanism identified here likely carries through to this more general setting as well.

## 3 Inconsistency of the Minimum Flip-Cost Perfect Phylogeny Objective

We now ask whether the MFCP objective can recover the true tree in principle. Specifically, does the true tree uniquely minimize the expected flip cost induced by the generative process?

### 3.1 Inconsistency

Because the total flip cost is additive across mutations and the assignment of each mutation can be optimized independently, the expected flip cost over a full matrix 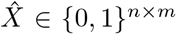 decomposes as a sum of independent per-mutation contributions as in Equation (1). It therefore suffices to analyze the expected cost of a single mutation. Let *ϕ*^∗^ denote the true edge on which the mutation occurs, i.e., for mutation *j, ϕ*^∗^ = Φ^∗^(*j*). We denote the expected flip cost of a candidate tree *T* under generative model (*T* ^∗^, **p**) by

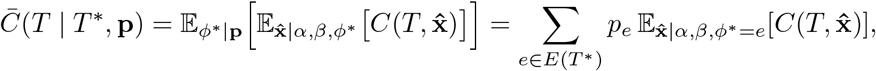

where the expectation is over both the edge *ϕ*^∗^ on which the mutation arises in *T* ^∗^ (conditional on fixed edge lengths **p**), and the observation noise generating 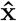 (conditional on *ϕ*^∗^ and the error probabilities *α* and *β*). In the following analyses we treat *α* and *β* as fixed parameters of the generative model and, for simplicity, suppress them from the notation of 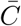 and all quantities derived from it.

#### ▶ Definition 5.

**Consistency**. *Consider a true tree T* ^∗^, *edge length distribution* **p** *on E*(*T* ^∗^), *and error rates α, β. The MFCP objective with weights w*_*N*_ , *w*_*P*_ *is* consistent *if the true tree T* ^∗^ *uniquely minimizes the expected cost:*

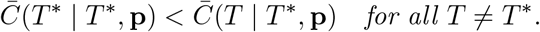

#### ▶ Remark 6.

This definition corresponds to Fisher consistency [21]. In our setting (where data are i.i.d., costs are bounded, and the tree space is finite) Fisher consistency is necessary and sufficient for asymptotic consistency, meaning the estimator would converge to the true tree as the number of mutations increases [46]. See Appendix A.1 for a detailed.

For our main result we will show that under the generative model defined in Section 2.1, consistency fails for all objectives in the MFCP family:

#### ▶ Theorem 7.

**Inconsistency of MFCP**. *For any weights w*_*N*_ , *w*_*P*_ *>* 0 *and any error probabilities α, β* ∈ (0, 0.5), *there exist a true tree T* ^∗^ *and an edge length distribution* **p** *for which the MFCP objective is not consistent*.

#### Intuition: structural absorption of errors

The key mechanism underlying inconsistency for this problem is a form of overfitting to observation noise. The MFCP objective will always assign mutations to whichever edge explains their errors most cheaply. As a result, the true number of observation errors is only an upper bound on the overall cost: a tree can achieve lower cost by “absorbing” some errors, reinterpreting a noisy pattern as a clean signal from a different edge. Whether a tree can absorb a given error depends on its topology.

As an example, consider the trees shown in Figure 1: a balanced tree ((*A, B*), (*C, D*)) and a caterpillar tree (((*A, B*), *C*), *D*) on four leaves, with a mutation arising on the trunk. This mutation has true pattern **x** = [1 1 1 1] (where the order of leaves is *A, B, C, D*). If a single false negative occurs on leaf *D*, this yields observed pattern 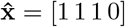. In the balanced tree the minimum cost assignment would require one flip (a 0 → 1 flip to the trunk [1 1 1 1], or a 1 → 0 flip to internal edge [1 1 0 0]) concordant with the true number of errors (Fig. 1A). In contrast, in the caterpillar tree, the same false negative produces an observed pattern that exactly matches the descendant set of the internal edge [1 1 1 0] (Fig. 1B). This allows the mutation to be assigned to the tree with zero flips, despite the fact that one observation error has occurred. Conversely, the balanced tree can explain two simultaneous false negatives on *A* and *B* with zero flips, while the caterpillar would require one flip. However, when false negative rate *α <* 0.5, single false negatives are more likely than double false negatives, so the caterpillar has a systematic advantage over the balanced tree for mutations arising on the trunk edge. We will show that these types of asymmetries are pervasive and unlikely to cancel out across edges.

**Figure 1.**
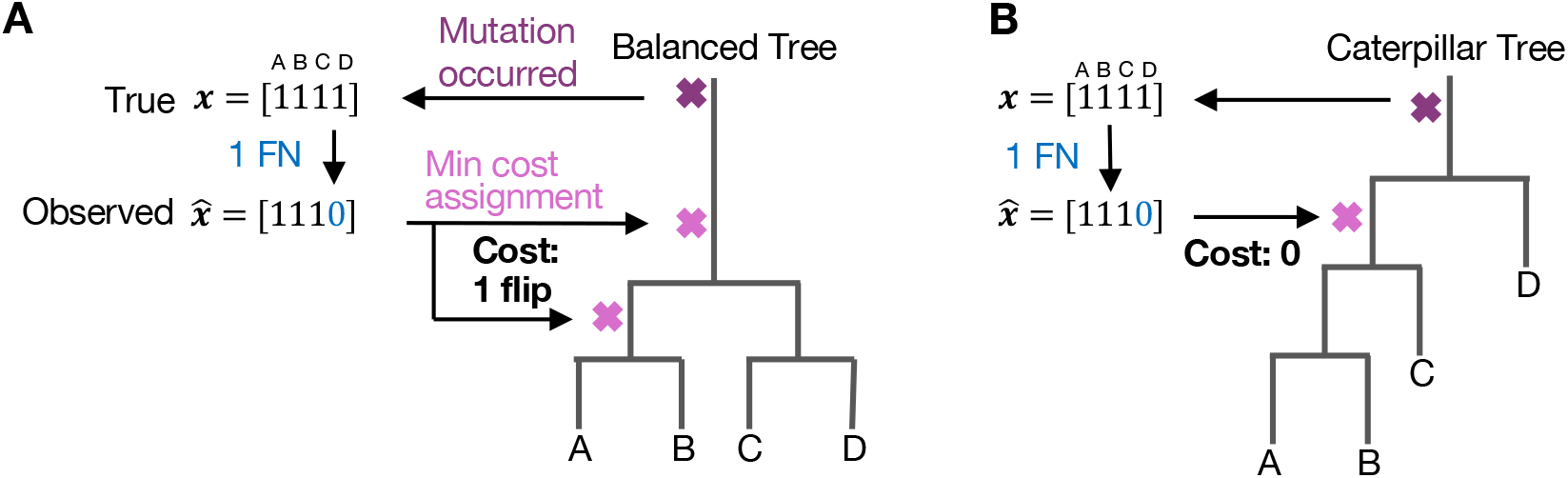
Structural absorption of observation errors. (**A**) A mutation arising on the trunk of a tree with balanced topology ((*A, B*), (*C, D*)) has true pattern **x** = [1 1 1 1], where the order of leaves is *A, B, C, D*. A false negative at leaf *D* produces observed pattern 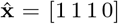, which is assigned to the minimum cost edge incurring a cost of one flip. (**B**) The same trunk mutation and false negative on the caterpillar tree (((*A, B*), *C*), *D*) produces the same observed pattern 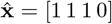. This observed pattern exactly matches the descendant set of the internal edge {*A, B, C*}. Therefore the minimum cost assignment for this pattern incurs zero cost, despite the fact that one observation error has occurred.

### 3.2 Per-edge cost decomposition

We first develop the machinery required to analyze the difference in expected cost between the true tree *T* ^∗^ and a candidate tree *T* . We decompose the expected cost difference Δ_**p**_(*T, T* ^∗^) into per-edge contributions:

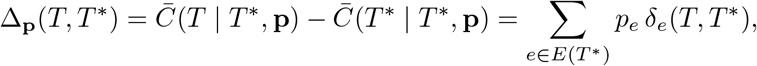

where *δ*_*e*_(*T, T* ^∗^) is the expected cost difference conditional on a mutation arising on edge *e* of *T* ^∗^:

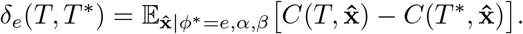

This decomposition allows us to analyze expected cost differences one generating edge at a time, separately from the edge length distribution. The following lemma shows that for consistency to hold, we would need *δ*_*e*_(*T, T* ^∗^) ≥ 0 for every *e* ∈ *E*(*T* ^∗^) and every *T T* ^∗^, as a single edge favoring a candidate tree is enough to break consistency (Fig. 2).

**Figure 2.**
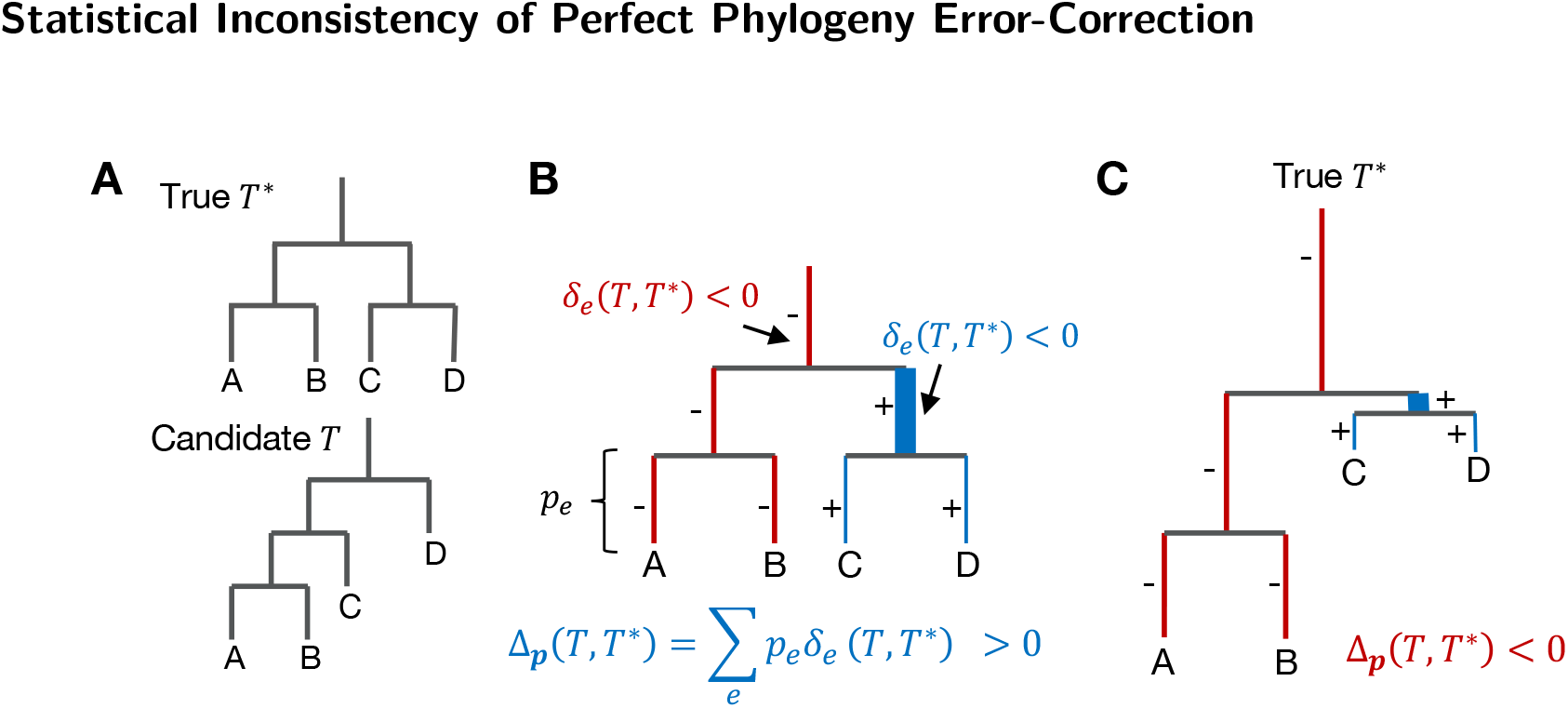
Edge-level cost differences and the edge concentration argument. (**A**) The true tree *T*^∗^ (balanced) and a candidate tree *T* (caterpillar). (**B**) The true tree *T*^∗^ with a balanced edge length distribution. The expected cost difference Δ_**p**_(*T, T*^∗^) between the candidate tree *T* and the true tree *T*^∗^ decomposes as a weighted sum of per-edge contributions *δ*_*e*_(*T, T*^∗^). Edge color indicates the sign of *δ*_*e*_(*T, T*^∗^) (red: negative, favoring the candidate; blue: positive, favoring the true tree), and edge width indicates magnitude. Under the balanced distribution, the weighted sum Δ_**p**_(*T, T*^∗^) is positive. (**C**) The true tree *T*^∗^ with a skewed edge length distribution that concentrates mass on the negatively-contributing edges. The sign and magnitude of *δ*_*e*_(*T, T*^∗^) remain unchanged from (**B**). However, because more mass is placed on the negatively-contributing edges, the weighted sum becomes negative.

#### ▶ Lemma 8.

**Edge concentration**. *Consider a true tree T* ^∗^ *and a candidate tree T* . *If there exists an edge e* ∈ *E*(*T* ^∗^) *such that δ*_*e*_(*T, T* ^∗^) *<* 0, *then there exists an edge length distribution* **p** *under which* 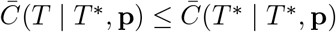.

**Proof**. Let *S*^−^ = {*e*^*′*^ ∈ *E*(*T* ^∗^) : *δ*_*e*_*′* (*T, T* ^∗^) < 0} be the set of edges favoring the candidate tree, and let *S*^+^ = {*e*^*′*^ ∈ *E*(*T* ^∗^) : *δ*_*e*_*′* (*T, T* ^∗^) ≥ 0} be the set of edges favoring the true tree. The expected cost difference can be expressed as the sum of contributions from the two sets ∑ _*e*_*′* _∈*S*_ − *p*_*e*_*′ δ*_*e*_*′* (*T, T* ^∗^) and ∑ _*e*_*′* _∈*S*_ + *p*_*e*_*′ δ*_*e*_*′* (*T, T* ^∗^). Since *S*^−^ is non-empty, the sum can be made non-positive by choosing **p** to concentrate sufficient mass on edges in *S*^−^ and away from edges in *S*^+^. ◀

If *e* is a shared split between two trees *T*_1_ and *T*_2_, then *δ*_*e*_(*T*_1_, *T*_2_) = −*δ*_*e*_(*T*_2_, *T*_1_). As a result, showing a single shared edge *e* such that *δ*_*e*_(*T*_1_, *T*_2_) ≠ 0 for some pair of trees *T*_1_ and *T*_2_ is sufficient to conclude that consistency is violated, by Lemma 8.

Lemma 8 is the device we use to establish inconsistency in the proofs that follow, by demonstrating examples of trees and edges for which *δ*_*e*_(*T, T* ^∗^) *<* 0. It assumes that edge lengths are unconstrained, so that mass can be redistributed toward the set *S*^−^ of negatively-contributing edges. This may raise concerns about whether violations of consistency require specially constructed or pathological edge length distributions. In Section 4, we will show that inconsistency does arise under reasonable edge length distributions and even for equal edge lengths.

### 3.3 Inconsistency of MFCP: a three-leaf counterexample

We provide two complementary proofs. The first is a concrete three-leaf example where the calculations can be confirmed directly; the second is a general construction for any tree topology with *n* ≥ 3 leaves, establishing that inconsistency is pervasive.

#### ▶ Lemma 9.

**Counterexample with three leaves**. *Let T and T* ^∗^ *be two distinct rooted binary trees on the same leaf set* {*A, B, C*}. *For any weights w*_*N*_ , *w*_*P*_ *>* 0 *and any error probabilities α, β* ∈ (0, 0.5), *there exists a probability distribution* **p** *such that* 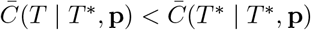.

**Proof**. Without loss of generality, label the leaves on the trees so that *T* ^∗^ = ((*A, B*), *C*) and *T* = (*A*, (*B, C*)), as shown in Figure 3A–B. We compute the expected cost difference 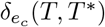 for the generating edge *e*_*c*_ corresponding to the outgroup leaf *C*, where 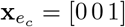. As shown in Figure 3C, among all eight possible observed patterns, the only ones for which *T* ^∗^ and *T* differ in cost are the two tree-specific internal splits [1 1 0] and [0 1 1]: each has cost 0 on the tree that contains it and cost one flip (i.e., min(*w*_*P*_ , *w*_*N*_)) on the other. The expected cost difference 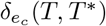 is therefore determined entirely by the relative probabilities of these two patterns: 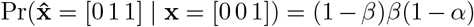 and 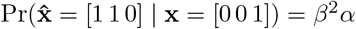. When *α, β* ∈ (0, 0.5), (1 − *β*)*β*(1 − *α*) *> β*^2^*α*, so 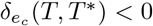. By Lemma 8, there exists a choice of edge lengths **p** such that 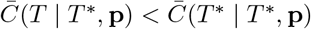. ◀

**Figure 3.**
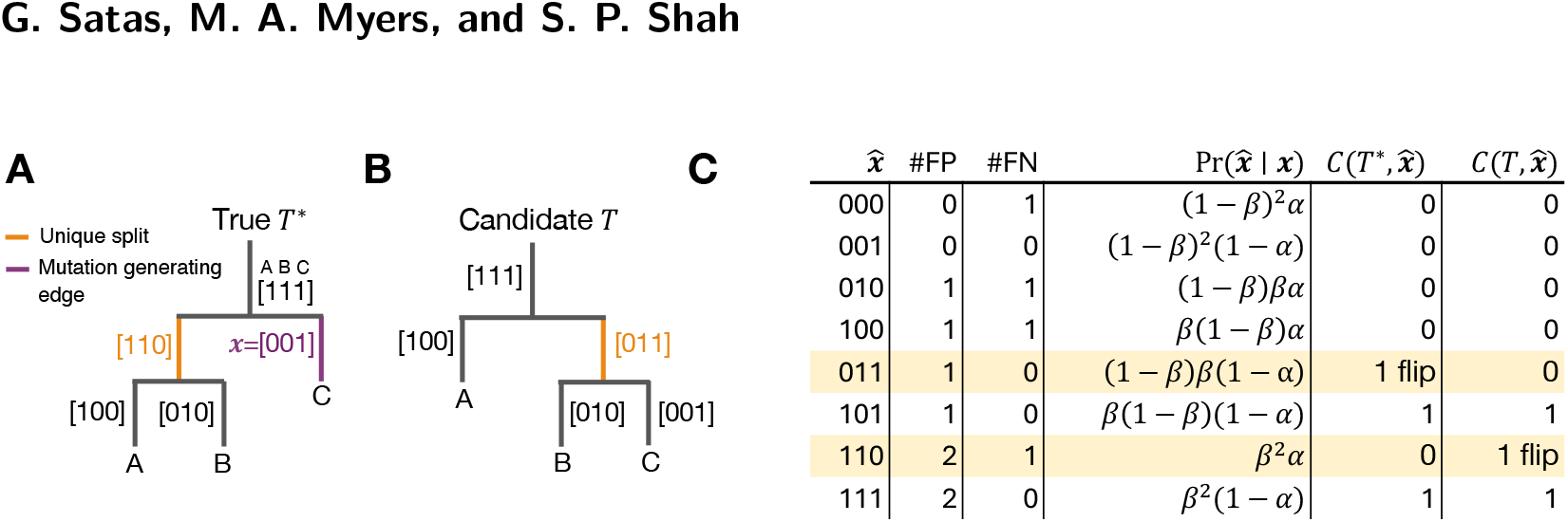
Three leaf violation of consistency (Lemma 9). (**A**) True tree *T*^∗^ on leaves *A, B, C*, with *A* and *B* as siblings. The split [1 1 0] (orange) is unique to *T*^∗^, where the order of leaves is *A, B, C*. The mutation-generating edge (purple) is the leaf edge of *C* with mutation pattern **x** = [0 0 1]. (**B**) Candidate tree *T* on the same leaf set, with *B* and *C* as siblings. The split [0 1 1] (orange) is unique to *T* . (**C**) All eight possible observed patterns 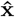 given true mutation state **x** = [0 0 1], with the number of false positives (#FP) and false negatives (#FN), the probability 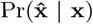, and the minimum flip cost 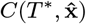 and 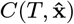 on each tree. The only patterns for which the two trees differ in cost are 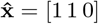 and [0 1 1] (orange).

Given Lemma 9, we can now prove the main Theorem 7.

#### Proof. Theorem 7.

For any weights and error probabilities, Lemma 9 provides a pair of distinct trees and an edge length distribution under which 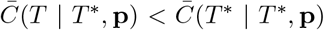.

Thus, by Definition 5, the MFCP objective is not consistent. ◀

### 3.4 Inconsistency of MFCP: a general construction for counterexamples

We now generalize the three-leaf counterexample to arbitrary tree topologies with *n* ≥ 3 leaves. We use pairs of trees that differ in exactly one internal split. These can be constructed for any tree with *n* ≥ 3 by performing a single rooted nearest-neighbor interchange (NNI) operation which swaps two subtrees across an internal edge [19]. Trees that differ by a single NNI are referred to as *NNI neighbors*. Any tree has two neighbors for every internal edge.

#### ▶ Lemma 10.

**Counterexample with NNI neighbors of arbitrary size**. *Let T*_1_ *and T*_2_ *be two distinct rooted binary trees on the same leaf set that are NNI neighbors. For any w*_*P*_ , *w*_*N*_ *>* 0 *and any α, β* ∈ (0, 0.5), *there exists an assignment of T*_1_ *and T*_2_ *to the roles of true tree T* ^∗^ *and candidate tree T* , *and an edge length distribution* **p** *such that* 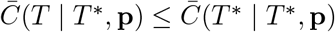.

We provide a full proof in Appendix A.2, but we give a sketch of the main idea here. An edge *e* that is shared between the two trees may be more similar to the unique split of a candidate tree *T* than that of the true tree *T* ^∗^, such that the edge is more likely to generate patterns that are assigned to the unique split of *T* than *T* ^∗^ at minimum cost. Therefore, for that edge, the expected cost of *T* is lower than that of *T* ^∗^. The three-leaf example is a minimal case demonstrating this, where the generating edge *e*_*c*_ = [0 0 1] was strictly more similar to the unique split [0 1 1] of *T* than [1 1 0] of *T* ^∗^. The proof in the Appendix guarantees that at least one such edge *e* exists for any pair of neighbors *T*_1_, *T*_2_ such that *δ*_*e*_(*T*_1_, *T*_2_) 0.

Lemma 10 provides not just a generic statement of existence, but also the basis for a constructive procedure for generating explicit counterexamples for any pair of neighbor trees: Given *any* neighbor pair (*T*_1_, *T*_2_), an inconsistent edge distribution can be found by (a) computing all *δ*_*e*_ values by enumerating the 2^*n*^ observed patterns, and (b) solving for an edge distribution **p** such that ∑_*e*_ *p*_*e*_*δ*_*e*_ *<* 0. Full details of the procedure are given in Appendix B.1, and an implementation that generates random examples of consistency violations under different edge length priors is available at https://github.com/shahcompbio/phylo_inconsistency.

### 3.5 The estimator can be positively misleading

Inconsistency, as established above, means only that the true tree fails to be the unique minimizer. A more severe form is when having more data is actively counterproductive, because the estimator converges to a specific incorrect tree with increasing data, known as estimator being *positively misleading* [18].

#### ▶ Definition 11.

**Positively misleading**. *The MFCP objective is* positively misleading *if the estimator converges to a unique tree T* ≠ *T* ^∗^ *rather than the true tree T* ^∗^ *as the number of mutations increases*.

#### ▶ Proposition 12.

**The MFCP is positively misleading**. *For any weights w*_*N*_ , *w*_*P*_ *>* 0 *and any error probabilities α, β* ∈ (0, 0.5), *there exist a true tree T* ^∗^ *and an edge length distribution* **p** *for which the MFCP objective is positively misleading*.

The proof of Proposition 12 is given in Appendix A.3. Briefly, it uses the counterexample from Lemma 9, with a constructed **p** to ensure that the minimizing tree is unique.

### 3.6 Inconsistency extends beyond nearest neighbors

Our counterexamples so far use NNI neighbors, but the phenomenon is far more general. By Lemma 8, a pair of trees *T*_1_, *T*_2_ can escape inconsistency only if *δ*_*e*_(*T*_1_, *T*_2_) = 0 on *every* shared edge. Each such expectation is a piecewise polynomial in the parameters (*α, β, w*_*P*_ , *w*_*N*_), and the condition *δ*_*e*_(*T*_1_, *T*_2_) = 0 for all *e* is an over-determined system of equations (at least *n* + 1 equations in three free parameters). Thus, the exact, simultaneous vanishing of the per-edge cost expectations across all shared edges, in general, fails for all but a measure-zero set of parameters (*α, β, w*_*P*_ , *w*_*N*_).

An exception to this is a structurally special family of pairs of trees whose shared-edge cost functionals cancel identically, so that *δ*_*e*_(*T*_1_, *T*_2_) = 0 for every shared edge *e*. One such example is the balanced quartet ((1, 2), (3, 4)) versus ((1, 3), (2, 4)). We do not fully characterize this family, but note that at minimum, the pair of trees must be isomorphic (although this is not a sufficient condition, see *n* = 3 counterexample). Thus the proportion of such pairs is relatively small, especially as *n* grows.

Consequently, guaranteed consistency is highly constrained and fragile in tree space, and counterexamples are available for nearly all pairs of trees and nearly all parameter values.

## 4 The prevalence of inconsistency under reasonable model parameters

### 4.1 Equal edge lengths do not guarantee consistency

Equal edge lengths is the most favorable candidate for consistency. Under the edge concentration argument, inconsistency requires only that the set *S*^−^ = {*e* : *δ*_*e*_ < 0} is non-empty; arbitrary edge lengths allow mass to be redistributed toward *S*^−^ and away from positively-contributing edges, making the negative contributions dominate. Equal edge lengths removes this freedom: the weighted sum Δ_**p**_ = ∑_*e*_ *p*_*e*_*δ*_*e*_ becomes a simple average, so inconsistency can only occur if the negative-*δ*_*e*_ edges outweigh the positive ones across the entire tree, a much stricter condition.

For *n* ≥ 4, we demonstrate computationally that consistency can still fail for some pairs of trees under equal edge lengths, and that the space of error parameters where this occurs is non-trivial and grows with *n*. We use three trees at each *n* (Figure 4A): *T*_1_ (full caterpillar), *T*_2_ (root has a 2-leaf right child), and *T*_3_ (root has a 3-leaf right child), where each tree is a neighbor of the previous one. For each tree size *n* ∈ [4, 9] (for *T*_1_/*T*_2_) and *n* ∈ [5, 9] (for *T*_2_/*T*_3_), we computed all four pairwise cost differences under equal edge lengths across a 25 *×* 25 grid of error probabilities (*α, β*), with weights matched to the error rates. We show that inconsistency occurs in all four directions, but asymmetrically (Figure 4B,C). *T*_2_ beats *T*_1_ over a substantial and growing region of parameter space, while *T*_1_ beats *T*_2_ only marginally: at *n* = 4, *T*_1_ is preferred over *T*_2_ for approximately 1% of parameter values, and this advantage disappears at larger *n*. Similarly, *T*_3_ beats *T*_2_ over a non-trivial region, while *T*_2_ beats *T*_3_ only at *n* = 6 for approximately 4% of parameter values. Inconsistency in general is concentrated at high error rates, and the space of error parameters where violations occur grows with *n* (Figure 4C); at *n* = 9 more than 50% of parameter values produce inconsistency in at least one direction, and the space does not appear to be saturating. This result is still weaker than the theoretical results, which cover *all* error parameters; however, these analyses were limited to a small set of tree topologies and tree sizes. It thus remains open whether any error regime fully restores consistency under equal edge lengths for trees of arbitrary size.

**Figure 4.**
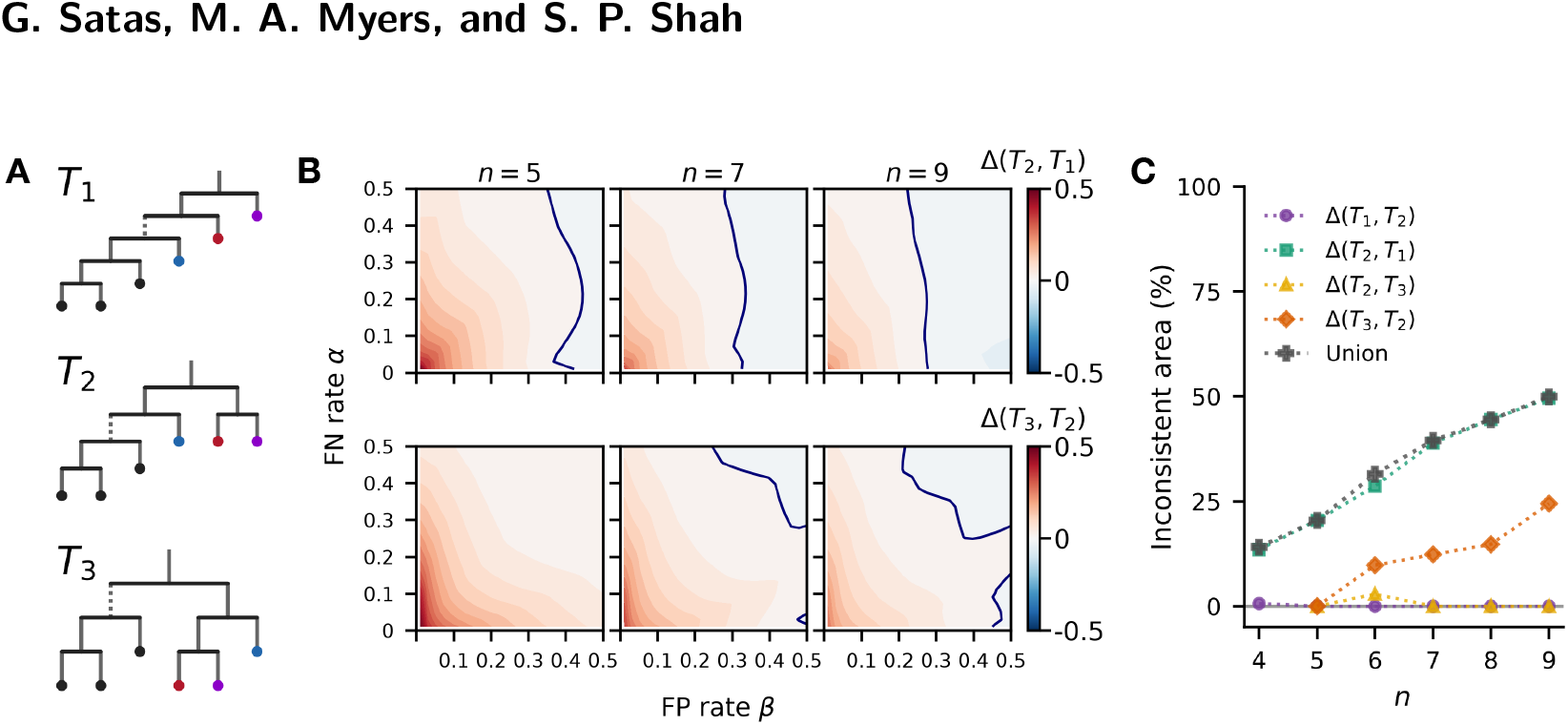
Consistency violations under equal edge lengths. (**A**) Three tree topologies on *n* leaves, named by the number of leaves on the right child of the root: *T*_1_, *T*_2_ and *T*_3_. Colored leaves are shared across all three trees, indicating which leaves change position. Increasing *n* adds leaves to the left subtree. (**B**) Heatmaps of the expected cost difference over a grid of error probabilities (*α, β*), shown for *n* ∈ {5, 7, 9} leaves. Top row: Δ_**p**_(*T*_2_, *T*_1_). Bottom row: Δ_**p**_(*T*_3_, *T*_2_). The blue line separates positive and negative regions, where the candidate has lower expected cost than the true tree. (**C**) Area of the inconsistent region as a percentage of the *α*-*β* parameter space, across all tested *n*. Each line corresponds to one direction of comparison: Δ_**p**_(*T*_2_, *T*_1_) *<* 0 (purple), Δ_**p**_(*T*_1_, *T*_2_) *<* 0 (green), Δ_**p**_(*T*_3_, *T*_2_) *<* 0 (amber), Δ_**p**_(*T*_2_, *T*_3_) *<* 0 (orange); black: union across all four directions.

### 4.2 Inconsistency is common under non-pathological edge-length distributions

To evaluate how often such counterexamples occur under biologically motivated edge length distributions, we simulated trees and edge length distributions using a Dirichlet distribution under a set of four different models determining parameters 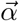: (a) *Balanced*: balanced edge length distribution, (b) *Non-informative*: uniform on the unit simplex, (c) *Heavy Trunk*: a long trunk with uniform internal edges, and (d) *Clade-weighted*: *p*_*e*_ ∝ |*D*_*e*_|^2^ (Figure 5A). Trees were generated with *n* ∈ [5, 9] leaves, with the maximum *n* limited by the computational cost of enumerating all 2^*n*^ observed patterns for each tree-edge length combination. For each *n* value, we simulated 500 trees, and for each tree, we took 1000 draws of edge length distributions under each model, for a total of 2,000,000 tree-edge length-model combinations. For each simulated tree and edge length distribution, we computed Δ_**p**_(*T, T* ^∗^) for all NNI neighbors *T* of the true tree *T* ^∗^. NNI neighbors were chosen as a relevant set of candidate trees as we expect them to be most likely to be preferred over the true tree, and the number of NNI neighbors scales linearly with *n*, making the analysis tractable. For further details on the simulation procedures, see Appendix B.2.

**Figure 5.**
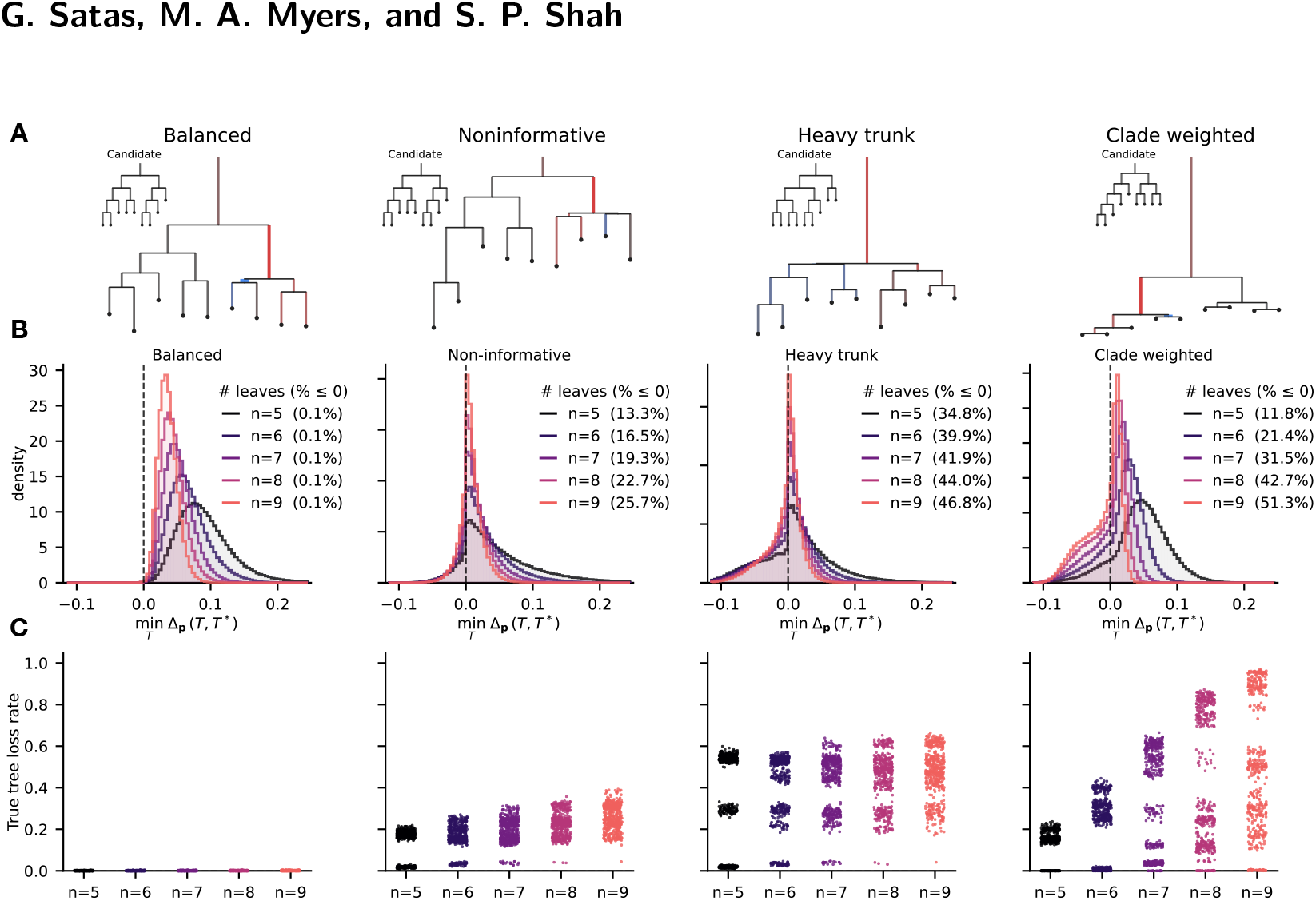
Prevalence of inconsistency under random edge length draws. Simulation results for trees with *n* leaves under four different edge length priors (left to right): balanced, non-informative, heavy trunk, and clade-weighted. (**A**) An example tree topology with *n* = 9 leaves. Each panel shows the same topology under an edge length distribution from each prior, where a candidate tree (shown as an inset) beats the true tree. Edge color indicates the sign of *δ*_*e*_(*T, T*^∗^) relative to the winning candidate shown in the inset (red: negative, favoring the candidate; blue: positive, favoring the true tree), and edge width indicates magnitude. Edge length is proportional to *p*_*e*_. (**B**) 500 trees are simulated for each *n* ∈ [5, 9], and for each tree, 1000 edge length draws are taken under each model. For each tree and edge length draw, we compute the minimum margin min_*T*_ Δ_**p**_(*T, T*^∗^) across all neighbors *T* of the true tree *T*^∗^, which is negative if at least one neighbor has a lower cost. For each model, the distribution of minimum margins is shown with the fraction of draws with a negative minimum margin in parentheses. (**C**) The distribution of true tree *loss rates*, i.e., the fraction of draws under which at least one neighbor beats the true tree. Each point corresponds to a single tree and shows the average loss rate across all draws.

For each tree-edge length combination, we computed the minimum margin min_*T*_ Δ_**p**_(*T, T* ^∗^) across all neighbors *T* of the true tree *T* ^∗^, which is negative if at least one neighbor achieves lower expected cost than the true tree under that edge length distribution. As expected, balanced edge lengths perform best: across all *n*, only ≈ 0.1% of draws produced any inconsistent neighbor (Figure 5B). Under the non-informative prior, inconsistency is substantially more common, and increases with *n*: at *n* = 5, 13.3% of draws produced at least one inconsistent neighbor, while at *n* = 9, this fraction increased to 25.7% (Figure 5B). The heavy trunk and clade-weighted priors also show increasing inconsistency with *n*, but at higher rates: 34.8% and 11.8% at *n* = 5 under the heavy trunk and clade-weighted priors respectively, increasing to 46.8% and 51.3% at *n* = 9 (Figure 5B).

The distributions show qualitatively distinct shapes across models (Figure 5B), with the heavy-trunk and clade-weighted models showing a pronounced concentration of mass in the negative range that increases with *n*. Notably, except under the balanced model, even when the true tree is preferred over all neighbors, the margin is often small, suggesting that in finite samples the true tree may frequently be indistinguishable from its neighbors. Moreover, at larger *n* under the heavy-trunk and clade-weighted models, the negative tail of the distribution extends further than the positive tail, indicating that when a neighbor beats the true tree, it often does so by a larger margin than the true tree achieves when it wins.

### 4.3 Susceptibility to inconsistency is dependent on tree topology and edge length distribution

We next examined the heterogeneity across topologies in their susceptibility to inconsistency. For each tree with *n* = 9, we computed the *true tree loss rate*: the fraction of the 1000 edge length draws under which at least one neighbor beats the true tree (Figure 5C and Figure A1A–C). Loss rates grow with the informativeness of the prior: nearly all trees stay below 0.5% under balanced edge lengths but reach medians of 0.248, 0.464, and 0.503 under the non-informative, heavy-trunk, and clade-weighted priors respectively, with the clade-weighted prior producing the widest spread (from near 0 to near 1). Under every structured prior essentially no tree is immune, and even under balanced edge lengths only 40% of trees never lost. The spread of loss rates widens with *n*, as some topologies become more susceptible and others more robust as tree space grows.

Strikingly, the topologies most susceptible under one prior are largely uncorrelated with those most susceptible under another (Figure A1A–C): trees form well-separated clusters in pairwise loss-rate space that subdivide further at larger *n*. Susceptibility is therefore not a fixed property of topology but is jointly determined by the tree and the edge-length distribution, with different priors placing the burden of inconsistency on different regions of tree space. One exception is the caterpillar topology, which consistently achieves among the lowest loss rates across all priors and tree sizes (Section 4.4).

The clustering suggests that loss rates are driven by discrete topological features that differ across priors. We fit random forest models predicting per-tree loss rate from eight topological features under each prior, excluding the balanced model, whose loss rates showed no meaningful variation across trees (see Appendix B.2.1 for details). Predictive accuracy was high in all three cases (*R*^2^ = 0.93, 0.97, and 0.98 for the non-informative, heavy-trunk, and clade-weighted priors; Figure A1D), confirming that susceptibility is systematically related to tree shape. The predictive features differed across priors in interpretable ways: global balance measures (e.g., number of cherries, Colless index) under the non-informative prior; root-proximal features (root split balance, whether the root has a singleton child) under the heavy-trunk prior, reflecting the influence of the trunk edge; and a mix of balance and depth features with non-monotonic effects under the clade-weighted prior, likely reflecting the dependence of its edge lengths on clade sizes. While the relationship between topology and susceptibility is complex and prior-dependent, identifiable structural properties of the tree thus interact with the edge-length distribution to determine which topologies are most at risk.

### 4.4 Bias toward caterpillar-like trees at realistic scDNA error rates

We examined which topological features systematically differ between neighbors that beat the true tree and the true tree itself (Figure 6A). These simulations use error rates calibrated to single-cell sequencing data (*α* = 0.2, *β* = 0.01), where false negatives are substantially more common than false positives. In this regime, winning neighbors consistently had higher cherry counts and lower Colless indices than the true tree across all tested tree sizes and edge-length priors, indicating that the favored topologies tend to be more caterpillar-like. This is consistent with the observation that the caterpillar topology has the lowest loss rate across all priors and tree sizes in this regime (Figure 5C).

**Figure 6.**
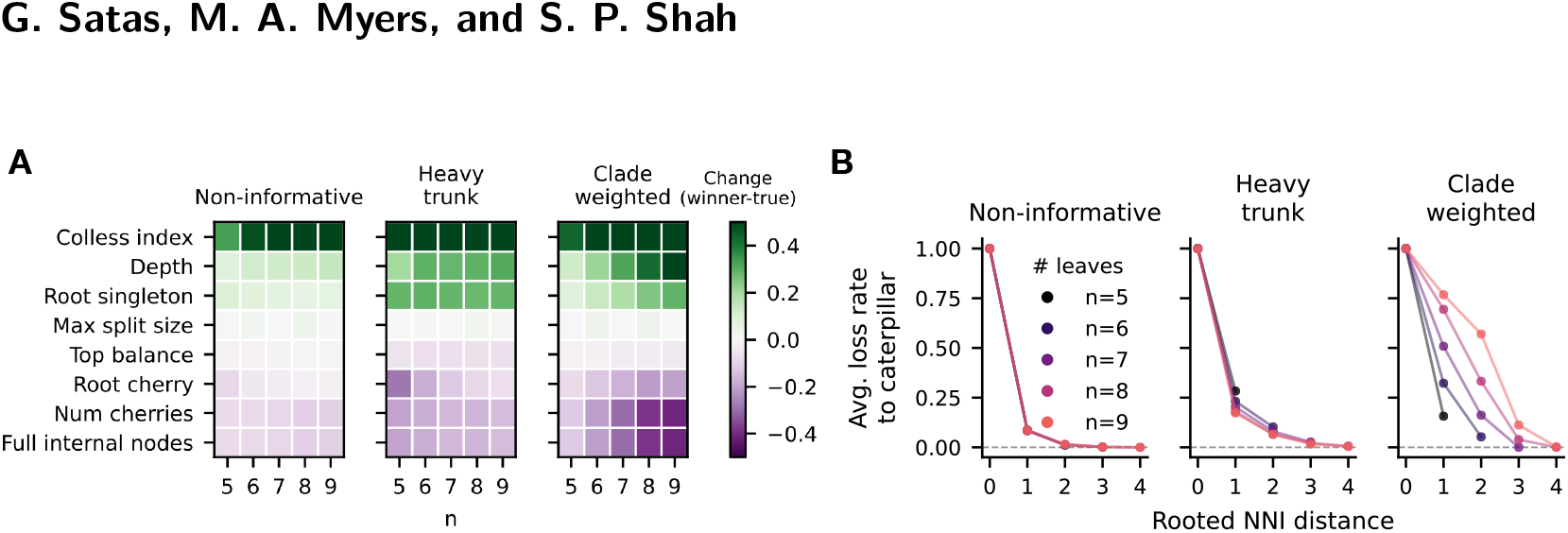
Topological bias toward caterpillar-like trees at realistic scDNA error rates. Simulations use error rates *α* = 0.2, *β* = 0.01 and three edge-length priors (Non-informative, Heavy trunk, Clade-weighted); the Balanced prior is omitted here as inconsistency is negligible in that regime. (**A**) Mean difference in topological features between winning neighbors and the true tree, averaged over all edge-length draws, across tree sizes *n* ∈ {5, … , 9} and the three priors. Positive values (red) indicate the winning neighbor tends to have a higher value of that feature than the true tree; negative (blue) indicates the reverse. (**B**) Rate at which a heuristic closest caterpillar beats the true tree in expected flip cost, as a function of the NNI distance between the caterpillar and the true tree. Uses the same error rates and edge-length priors as Panel A, but instead of comparing each true tree against all of its neighbors, each true tree is compared against a single fixed candidate: the heuristic closest caterpillar (see Appendix B.2.1), which may be more than one NNI move away. Each point is the fraction of edge-length draws where the caterpillar wins, averaged over all true trees at a given NNI distance, stratified by tree size and edge-length prior.

We note that this directional bias is specific to this error-rate regime and should not be taken as a general property of the inconsistency phenomenon. Under more symmetric or reversed error rates the preferred topologies shift. This is highly evident in the equal edge-lengths analysis (Section 4.1): at high false positive rates, the bias strongly favors the more balanced trees. Thus, the parameter dependence is not a fixed artifact of the objective but changes with the error profile of the data.

### 4.5 Inconsistency extends beyond neighboring tree topologies

In Section 3.6, we argued that inconsistency is not confined to NNI neighbors of the true tree, but can extend to more distant regions of tree space. We now provide empirical evidence that this is the case, by comparing the true tree against a candidate that is more topologically distant than its immediate neighbors. Because the caterpillar consistently achieves the lowest loss rate in this error regime, we used a heuristic closest caterpillar (see Appendix B.2.1 for construction details) as a fixed candidate and computed how often it beat the true tree as a function of NNI distance. Even when the candidate is several splits away from the true tree, it achieves lower expected cost in a non-trivial fraction of cases (Figure 6B). This is most pronounced under the clade-weighted prior, where the caterpillar beats the true tree in ≈ 50% of cases at an NNI distance of 2 and ≈ 15% at an NNI distance of 3 when *n* = 9, and the proportions increase with *n*. These results are a lower bound on the severity of long-range inconsistency: the caterpillar labeling used is a heuristic, and other labelings or topologically distant candidates may beat the true tree even more frequently, but were not evaluated due to computational constraints. Nonetheless, this suggests that inconsistency is not confined to small topological perturbations of the true tree but can extend to more distant regions of tree space.

## 5 Discussion

We studied the consistency of perfect phylogeny error-correction objectives under observation noise. Our central result is a general inconsistency phenomenon: hard-assignment objectives such as the minimum-cost perfect phylogeny (MFCP) objective and its variants are not guaranteed to uniquely favor the true evolutionary tree in expectation and the failure can be positively misleading: increasing the number of observed mutations can strengthen the support for an incorrect tree. This failure arises from the interaction between observation noise and the structure of the optimization objective itself. It can arise even under equal edge lengths, balanced trees, and symmetric error rates, and is therefore not a consequence of pathologically extreme parameter choices or topologies.

A key feature of the inconsistency phenomenon is that violations are systematic rather than random. Our computational results show that some topologies may be systematically favored across a wide range of edge length distributions. The pattern of which topologies are favored correlates with identifiable structural features, but we do not yet have a complete characterization of these features or how they interact with the observation rates (*α, β*) and edge-length distribution to produce bias. While we have shown that trees can be beaten by candidates that are several splits away, one important open question is a more complete characterization of how far the global optimum typically is from the true tree in practice.

If systematic biases operate across an entire cohort of samples, they may be misinterpreted as recurrent biological signal when they are in fact artifacts of the objective. The direction of bias depends on the observation error regime: under high false negative rates, as is common in single-cell sequencing, the bias favors more linear topologies; under high false positive rates, balanced topologies tend to be preferred instead. The distinction between branching and linear tumor phylogenies has been a topic of considerable interest in cancer genomics [12, 13], and some methods have used error-correction objectives specifically to ask whether a linear phylogeny is compatible with observed data [3, 48]. Understanding when and how biases in the objective operate is directly relevant for interpreting the results of such analyses.

We wish to emphasize that this paper is not a call to abandon these error-correction objectives in practice. Inconsistent estimators are widely used and often adequate for their intended purpose. While our results show that inconsistency can arise under biologically motivated edge lengths and error rates calibrated to scDNA sequencing, the extent to which it affects biological interpretation in real datasets remains an important direction for future work.

As a consistent alternative, we speculate that maximum-likelihood approaches that marginalize over mutation assignments and edge lengths may be less susceptible to this phenomenon, as they do not rely on a single hard assignment of observed patterns to tree splits. However, marginal approaches often carry additional costs: naive implementations of marginal likelihoods are computationally expensive as they require exhaustive enumeration of all values. Developing consistent and computationally practical alternatives to hard-assignment objectives, and characterizing the conditions under which they are needed, remain important open problems.

## A Proofs and derivations

### A.1 Fisher Consistency and Asymptotic Consistency

Remark 6 asserts that, under our generative model, Fisher consistency (Definition 5) is necessary and sufficient for asymptotic consistency. We justify this here, and more generally identify the tree to which the estimator converges. Consider a true tree *T* ^∗^, edge lengths **p**, and error rates *α, β*. For a sample of *m* i.i.d. mutations, let *Ĉ*_*m*_(*T*) denote the empirical flip cost of tree *T* and 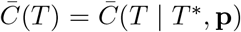 its expectation. The MFCP estimator selects 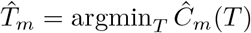.

Suppose the expected cost 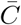 has a unique minimizer *T* ^min^ over the tree space, i.e. 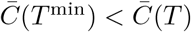 for all *T ≠ T* ^min^. Then 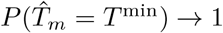 as *m* → ∞. This follows from Theorem 5.7 of [46], whose three conditions all hold here: (i) the mutations are i.i.d. by assumption; (ii) each per-mutation cost 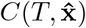 is bounded (at most *n* flips), so in particular has finite expectation; (iii) the tree space is finite, so 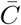 has a well-separated unique minimizer at *T* ^min^ (with 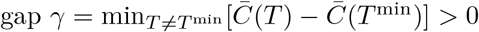).

When Fisher consistency holds, *T*^min^ = *T*\*, so 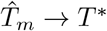 and the estimator is asymptotically consistent. Conversely, suppose Fisher consistency fails, so there exists some *T* ≠ *T* ^∗^ with 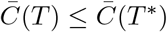. By the law of large numbers 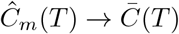 and 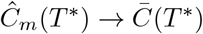, so *Ĉ*_*m*_(*T*) ≤ *Ĉ*_*m*_(*T* ^∗^) with probability approaching 1, and the estimator does not converge to *T* ^∗^.

### A.2 Proof of Lemma 10

We use the setup and notation of Lemma 10 throughout this section: *T*_1_ and *T*_2_ are NNI neighbors with subtrees *t*_*A*_, *t*_*B*_, *t*_*C*_, *t*_*D*_ where *t*_*A*_ and *t*_*B*_ are the exchanged subtrees. *T*_1_ has unique split *s*_1_, which is ancestral to subtrees *t*_*A*_ and *t*_*C*_, while *T*_2_ has unique split *s*_2_ which is ancestral to subtrees *t*_*B*_ and *t*_*C*_. Let *A* be a leaf of subtree *t*_*A*_, and *B* be a leaf of subtree *t*_*B*_. Without loss of generality, we order mutation patterns **x** = [*x*_*A*_ *x*_*B*_ **z**] where *x*_*A*_ and *x*_*B*_ are the state of *A* and *B* respectively and 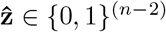 is the state of the remaining leaves.

Let *e*_*A*_ and *e*_*B*_ be the edges corresponding to leaves *A* and *B*. Below we will demonstrate the following

1. Patterns of the form 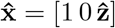 relatively benefit *T*_1_ when compared to patterns of the form 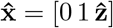.
2. Patterns of the form 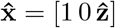 are more likely to be generated by *e*_*A*_ than patterns of the form 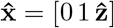.

By symmetry, the opposite of these properties hold for *T*_2_ and *e*_*B*_. As a result, *e*_*A*_ is more likely to generate patterns that benefit *T*_1_ while *e*_*B*_ is more likely to generate patterns that benefit *T*_2_, and consequently 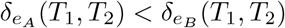.

#### Cost Asymmetry

Here we show that for all 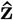, the net advantage 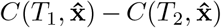 strictly decreases between observed pattern 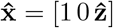 and 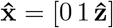. That is patterns of the form 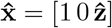 relatively benefit *T*_1_, compared to patterns of the form 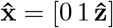 which relatively benefit *T*_2_.

Tree *T*_1_ can only achieve a lower cost on a pattern 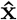 than *T*_2_ if the minimal cost assignment for 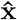 is the unique split *s*_1_, because *T*_2_ shares all other splits in *T*_1_. Let *n*_*A*_ be the number of leaves in *t*_*A*_, and let *k*_*A*_ be the number of leaves in *t*_*A*_ with state 1, for some 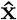.

*s*_1_ can only be the minimum cost assignment if *w*_*N*_ · *k*_*A*_ ≥ *w*_*P*_ · (*n*_*A*_ − *k*_*A*_); that is, the cost of flipping all 1 → 0 exceeds the cost of flipping all 0 → 1. Otherwise, the trunk of subtree *t*_*C*_ would have lower cost than *s*_1_. Note that this is a necessary but not sufficient condition for *s*_1_ to be the optimal assignment. Consider patterns of the form 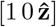 and 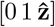 for 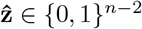. Switching from 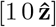 to 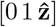 strictly decreases the number of leaves in *t*_*A*_ with value 1. Therefore, it can never make *s*_1_ optimal if it were not already optimal. A symmetric argument holds for *T*_2_ and flipping from 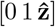 to 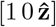. As a result, we have that

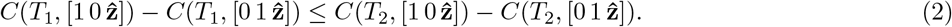

There exists at least one 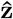 for which this inequality is strict. Take 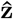 with all leaves of *t*_*B*_ and *t*_*D*_ in state 0 and all leaves of *t*_*C*_ in state 1, and choose the number of state-1 leaves in *t*_*A*_ so that the unique split *s*_1_ is the unique minimum-cost assignment for 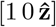 but not for 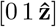. Such a choice exists: passing from 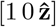 to 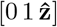 flips *A* from 1 to 0, removing one leaf from *t*_*A*_ and hence from the clade of *s*_1_, and there is a number of state-1 leaves in *t*_*A*_ at which this single flip moves *s*_1_ from uniquely optimal to non-optimal (tied or beaten by the *t*_*C*_ split). For this 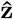, *T*_1_ attains via *s*_1_ a cost on 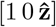 that *T*_2_ cannot match, whereas on 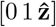 any cost *T*_1_ attains is also attained by *T*_2_ through a shared split. Therefore, 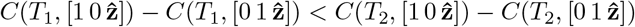.

#### Probability asymmetry

Under the generating edge *ϕ*^∗^ = *e*_*A*_, leaf *A* has true state 1 and leaf *B* has true state 0; under *ϕ*^∗^ = *e*_*B*_ the opposite holds. Let *γ* = (1 − *α*)(1 − *β*) − *αβ* and 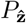 be the probability of observing 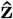. By independence of the observation model, the probabilities of each pattern type are:

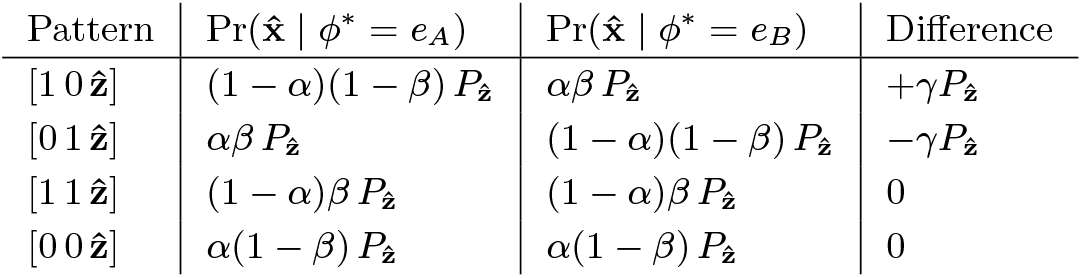

For all *α, β* ∈ (0, 0.5), *γ >* 0.

### A.3 Proof of Proposition 12

We use the three-leaf counterexample of Lemma 9. On leaf set {*A, B, C*} there are exactly three rooted binary trees,

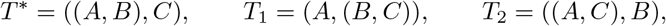

which share every split except their internal cherry: [1 1 0], [0 1 1], and [1 0 1] respectively. Note that candidates *T*_1_ and *T*_2_ are symmetric under the exchange of leaves *A* and *B* with respect to the true tree *T* ^∗^, and thus by the argument of Lemma 9, the generating edge *e*_*C*_ with true pattern [0 0 1] favors both *T*_1_ and *T*_2_ over *T* ^∗^ by the same amount. By the same argument, the generating leaf edges *e*_*A*_ and *e*_*B*_ with true patterns [1 0 0] and [0 1 0] favor *T*_2_ and *T*_1_ respectively. Consequently, making 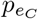 sufficiently large guarantees that both *T*_1_ and *T*_2_ beat *T* ^∗^, while making 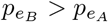 ensures that *T*_1_ beats *T*_2_. As a result, by Appendix A.1 the estimator uniquely converges to *T*_1_≠ *T* ^∗^ as the number of mutations increases.

### A.4 Derivation of Maximum Likelihood Weight Correspondence

We derive the flip-cost weights *w*_*P*_ and *w*_*N*_ under which the MFCP objective is equivalent to maximizing the joint likelihood over trees and mutation assignments. Consider a single mutation with observed pattern 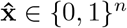. For each edge *e*, let **x** ∈ {0, 1}^*n*^ be the mutation pattern induced by *e*. The log-likelihood of assigning the mutation to edge *e* is

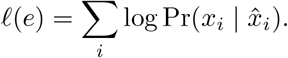

Let 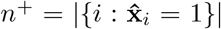 and 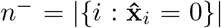, which depend only on 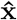 (not on *e*). The four cases for leaf *i* are:’

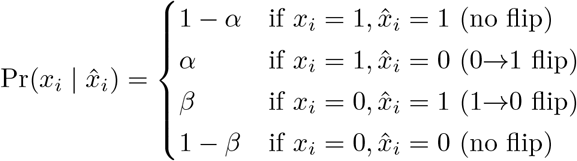

Let 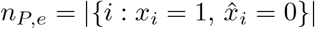 and 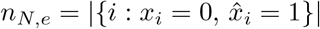. Collecting terms:

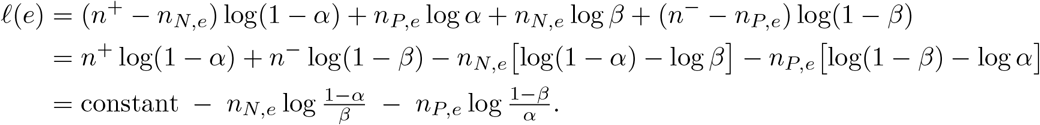

In other words, the log likelihood gives a penalty of log 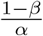 for each 0 → 1 flip and a penalty of 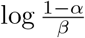 for each 1 → 0 flip, up to an additive constant that does not depend on the edge assignment. Summing over mutations and maximizing over (*T*, Φ) jointly therefore corresponds to the MFCP objective with weights 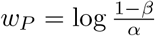 and 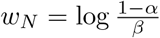.

## B Computational Methods

### B.1 Algorithmic Construction of Counterexamples

We describe a procedure for generating explicit counterexamples to consistency for any pair of neighbor trees, based on the proof of Lemma 10 and the edge concentration argument of Lemma 8. An implementation of this procedure is available at https://github.com/shahcompbio/phylo_inconsistency.

1. **Sample a random NNI pair**. Given *n* ≥ 3, we sample a random rooted binary tree *T* ^∗^ on *n* leaves. We then sample an internal edge uniformly at random and apply an NNI operation to obtain the candidate tree *T* .
2. **Compute** *δ*_*e*_. For each edge *e* ∈ *E*(*T* ^∗^) we compute *δ*_*e*_(*T, T* ^∗^) by explicit enumeration over all 2^*n*^ observed patterns 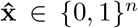. If all *δ*_*e*_ ≥ 0 (i.e. *T* ^∗^ already dominates pointwise), the roles of *T* ^∗^ and *T* are swapped, as Lemma 10 guarantees at least one orientation will have a negative delta.
3. **Find an inconsistent edge distribution**. We aim to find an edge distribution subject to *p δ <* 0 that is not pathologically extreme when possible. We look for the distribution that is closest (in KL divergence) to a chosen prior **q** subject to ∑_*e*_ *p*_*e*_*δ*_*e*_ = 0, then perturb it to make the sum strictly negative. The minimum-KL problem min KL(*p q*) subject to ∑_*e*_ *p*_*e*_*δ*_*e*_ = 0 and ∑_*e*_ *p*_*e*_ = 1 has a closed-form solution via Lagrange multipliers:

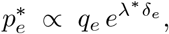

where *λ*^∗^ is the unique root of 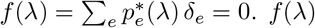 is continuous and monotone in *λ* so a root is found efficiently via binary search.

We use three priors **q**: (a) *Balanced:* all edges are equally weighted, (b) *Heavy trunk:* all edges are equally weighted except the trunk edge, which is given a higher weight, and (c) *Clade weighted:* 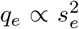, where *s*_*e*_ = ∑_*i*_ (**x**_*e*_)_*i*_ is the clade size of edge *e*. The clade-weighted prior is inspired by coalescent models of evolution, which predict that edges with larger clades tend to be longer.

### B.2 Simulation procedures

We describe the simulation pipeline used in Section 4.

#### Tree generation

For each simulation condition we draw *N* random rooted binary trees on *n* leaves.

#### Edge-length distributions

We treat the edge-length vector **p** = (*p*_*e*_)_*e E*(*T*_ ∗_)_, ∑_*e*_ *p*_*e*_ = 1, as a random quantity drawn from a prior distribution. We consider four priors.

1. **Balanced. p** ∼ Dirichlet(5).
2. **Non-informative. p** ∼ Dirichlet(**1**).
3. **Heavy-trunk. p** ∼ Dirichlet(***α***) where *α*_*e*_ = 5 for the trunk edge (the root edge) and *α*_*e*_ = 1 for all other edges.
4. **Clade-weighted. p** ∼ Dirichlet(***α***) where *α*_*e*_ = |*D*_*e*_|^2^.

#### NNI comparison

For each sampled true tree *T* ^∗^ and edge-length draw **p**, we test whether the objective prefers *T* ^∗^ over all of its NNI neighbours, i.e., Δ_**p**_(*T, T* ^∗^) *<* 0. For each internal edge *e* ∈ *E*(*T* ^∗^) and each of its two NNI rearrangements, we compute the margin

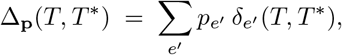

where *δ*_*e*_*′* (*T, T* ^∗^) is computed by explicit enumeration over all 2^*n*^ observed patterns 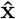.

#### B.2.1 True tree loss rate prediction

To investigate the relationship between tree topology and the loss rate of the true tree *T* ^∗^, we computed a set of topological features for each tree, as follows:

▄ **Depth**. The maximum distance from the root to any leaf.
▄ **Balance**. The Colless index, which is the sum over all internal nodes of the absolute difference in the number of descendant leaves in the left and right subtrees
▄ **Top balance**. The balance at the root node, i.e., the absolute difference in the number of descendant leaves in the left and right subtrees of the root.
▄ **Root has singleton child**. Whether the root has a child that is a leaf.
▄ **Root has cherry child**. Whether the root has a child that is an internal node with two leaf children (a cherry).
▄ **Cherry count**. The number of cherries in the tree, where a cherry is an internal node with two leaf children.
▄ **Full internal node count**. The number of internal nodes with two internal node children.
▄ **Max split size**. The size of the largest internal split of the tree, measured by the number of descendant leaves on the smaller side of the split.

A random forest regression model was then trained to predict the loss rate of the true tree *T* ^∗^ from these topological features, separately for each edge-length prior. The importance of each feature was evaluated using permutation importance, which measures the increase in mean squared error when the feature is randomly permuted across trees.

#### B.2.2 Heuristic Caterpillar Construction

There are many caterpillar trees on *n* leaves that differ in their leaf ordering along the spine. Some of these may be more likely to be preferred over the true tree than others, and it is unclear which caterpillar tree would be optimal. We used a heuristic construction to select a caterpillar by selecting the tree whose internal clades closely resemble the clades of the true tree. We extract a leaf ordering from the true tree through a depth-first traversal from the root. At each internal node, the deeper child clade is visited first. This construction is heuristic and doesn’t guarantee that the resulting caterpillar is the best possible caterpillar for comparison, but it is a principled way to select a single caterpillar that may be more likely to be preferred over the true tree than a random caterpillar, and it is computationally efficient to construct.

## C Additional Figures

**Figure A1.**
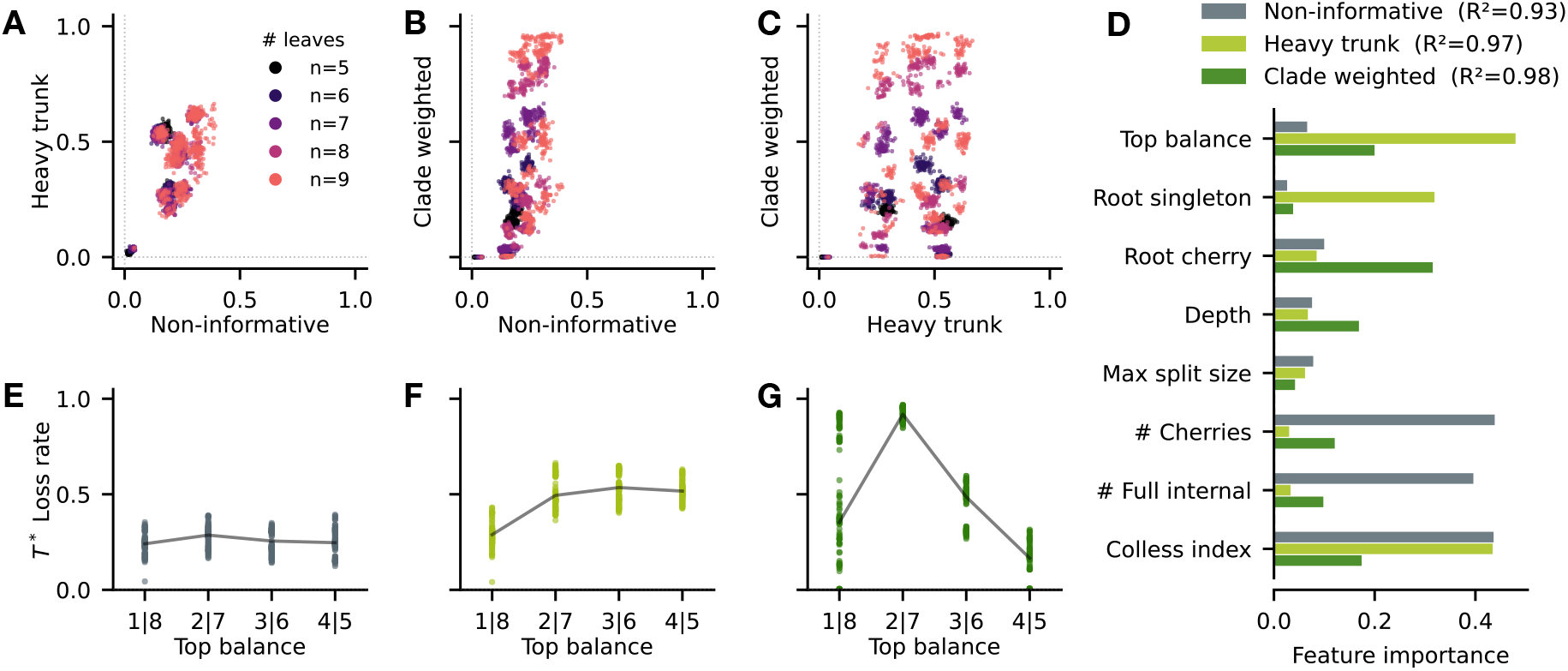
Prediction of true-tree loss rate from tree topology. (**A–C**) Pairwise scatter plots of true-tree loss rates across all tree topologies, comparing the Non-informative, Heavy trunk, and Clade-weighted edge-length priors. Each point represents one tree topology. (**D**) Permutation feature importance for the topological features used in the random forest models, shown for the three edge-length models with the same color scheme as in panels A–C: Non-informative in grey, Heavy trunk in orange, and Clade-weighted in purple. Cross-validated *R*^2^ values for each model are annotated. (**E–G**) Distributions of the top-balance feature for the trees under the Non-informative, Heavy trunk, and Clade-weighted models, respectively.

## References

1 Richa Agarwala and David Fernández-Baca. A polynomial-time algorithm for the perfect phylogeny problem when the number of character states is fixed. In Proceedings of 1993 IEEE 34th Annual Foundations of Computer Science, pages 140–147. IEEE, 1993.

2 Pranjal Awasthi, Avrim Blum, Jamie Morgenstern, and Or Sheffet. Additive approximation for near-perfect phylogeny construction. In International Workshop on Approximation Algorithms for Combinatorial Optimization, pages 25–36. Springer, 2012.

3 Erfan Sadeqi Azer, Mohammad Haghir Ebrahimabadi, Salem Malikić, Roni Khardon, and S Cenk Sahinalp. Tumor phylogeny topology inference via deep learning. Iscience, 23(11), 2020.

4 Niko Beerenwinkel, Roland F Schwarz, Moritz Gerstung, and Florian Markowetz. Cancer evolution: mathematical models and computational inference. Systematic biology, 64(1):e1–e25, 2015.

5 Sebastian Böcker, Quang Bao Anh Bui, François Nicolas, and Anke Truss. Intractability of the minimum-flip supertree problem and its variants. arXiv preprint arXiv:1112.4536, 2011.

6 Sebastian Böcker, Quang Bao Anh Bui, and Anke Truss. An improved fixed-parameter algorithm for minimum-flip consensus trees. In International Workshop on Parameterized and Exact Computation, pages 43–54. Springer, 2008.

7 John D Bridgers, Jan Hoinka, S Cenk Sahinalp, Salem Malikic, Teresa M Przytycka, and Funda Ergun. Improved algorithms for bi-partition function computation. In 25th International Conference on Algorithms for Bioinformatics (WABI 2025), pages 5–1. Schloss Dagstuhl–Leibniz-Zentrum für Informatik, 2025.

8 Malte Brinkmeyer, Thasso Griebel, and Sebastian Böcker. Flipcut supertrees: towards matrix representation accuracy in polynomial time. Algorithmica, 67(2):142–160, 2013.

9 Duhong Chen, Oliver Eulenstein, David Fernandez-Baca, and Michael Sanderson. Minimum-flip supertrees: complexity and algorithms. IEEE/ACM transactions on computational biology and bioinformatics, 3(2):165–173, 2006.

10 Markus Chimani, Sven Rahmann, and Sebastian Böcker. Exact ilp solutions for phylogenetic minimum flip problems. In Proceedings of the First ACM International Conference on Bioinformatics and Computational Biology, pages 147–153, 2010.

11 Simone Ciccolella, Camir Ricketts, Mauricio Soto Gomez, Murray Patterson, Dana Silverbush, Paola Bonizzoni, Iman Hajirasouliha, and Gianluca Della Vedova. Inferring cancer progression from single-cell sequencing while allowing mutation losses. Bioinformatics, 37(3):326–333, 2021.

12 Ibiayi Dagogo-Jack and Alice T Shaw. Tumour heterogeneity and resistance to cancer therapies. Nature reviews Clinical oncology, 15(2):81–94, 2018.

13 Alexander Davis, Ruli Gao, and Nicholas Navin. Tumor evolution: Linear, branching, neutral or punctuated? Biochimica et Biophysica Acta (BBA)-Reviews on Cancer, 1867(2):151–161, 2017.

14 James H Degnan, Michael DeGiorgio, David Bryant, and Noah A Rosenberg. Properties of consensus methods for inferring species trees from gene trees. Systematic Biology, 58(1):35–54, 2009.

15 James H Degnan and Noah A Rosenberg. Discordance of species trees with their most likely gene trees. PLoS genetics, 2(5):e68, 2006.

16 Mohammadamin Edrisi, Hamim Zafar, and Luay Nakhleh. A combinatorial approach for single-cell variant detection via phylogenetic inference. bioRxiv, page 693960, 2019.

17 Mohammed El-Kebir. Sphyr: tumor phylogeny estimation from single-cell sequencing data under loss and error. Bioinformatics, 34(17):i671–i679, 2018.

18 Joseph Felsenstein. Cases in which parsimony or compatibility methods will be positively misleading. Systematic zoology, pages 401–410, 1978.

19 Joseph Felsenstein. Inferring phylogenies. In Inferring phylogenies, pages 664–664. 2004.

20 Daniel W Feng and Mohammed El-Kebir. Dolphyin: A combinatorial algorithm for identifying 1-dollo phylogenies in cancer. In 25th International Conference on Algorithms for Bioinformatics (WABI 2025), pages 9–1. Schloss Dagstuhl–Leibniz-Zentrum für Informatik, 2025.

21 Ronald A Fisher. On the mathematical foundations of theoretical statistics. Philosophical transactions of the Royal Society of London. Series A, containing papers of a mathematical or physical character, 222(594–604):309–368, 1922.

22 Walter M Fitch. Toward defining the course of evolution: minimum change for a specific tree topology. Systematic Biology, 20(4):406–416, 1971.

23 Dan Gusfield. Efficient algorithms for inferring evolutionary trees. Networks, 21(1):19–28, 1991.

24 Michael D Hendy and David Penny. A framework for the quantitative study of evolutionary trees. Systematic zoology, 38(4):297–309, 1989.

25 Akhil Jakatdar, Palash Sashittal, and Benjamin J Raphael. Deriving longitudinal tumor phylogenies from single-cell sequencing data. bioRxiv, pages 2025–08, 2025.

26 Junhyong Kim. General inconsistency conditions for maximum parsimony: effects of branch lengths and increasing numbers of taxa. Systematic Biology, 45(3):363–374, 1996.

27 Motoo Kimura. The number of heterozygous nucleotide sites maintained in a finite population due to steady flux of mutations. Genetics, 61(4):893, 1969.

28 Laura Salter Kubatko and James H Degnan. Inconsistency of phylogenetic estimates from concatenated data under coalescence. Systematic biology, 56(1):17–24, 2007.

29 Liang Liu and Scott V Edwards. Phylogenetic analysis in the anomaly zone. Systematic biology, 58(4):452–460, 2009.

30 Adiesha Liyanage, Robyn Burger, Allison Shi, Braeden Sopp, Binhai Zhu, and Brendan Mumey. Essentcell: Discovering essential evolutionary relations in noisy single-cell data. bioRxiv, pages 2025–04, 2025.

31 Jian Ma, Aakrosh Ratan, Brian J Raney, Bernard B Suh, Webb Miller, and David Haussler. The infinite sites model of genome evolution. Proceedings of the National Academy of Sciences, 105(38):14254–14261, 2008.

32 Salem Malikic, Katharina Jahn, Jack Kuipers, S Cenk Sahinalp, and Niko Beerenwinkel. Integrative inference of subclonal tumour evolution from single-cell and bulk sequencing data. Nature communications, 10(1):2750, 2019.

33 Salem Malikic, Farid Rashidi Mehrabadi, Simone Ciccolella, Md Khaledur Rahman, Camir Ricketts, Ehsan Haghshenas, Daniel Seidman, Faraz Hach, Iman Hajirasouliha, and S Cenk Sahinalp. Phiscs: a combinatorial approach for subperfect tumor phylogeny reconstruction via integrative use of single-cell and bulk sequencing data. Genome research, 29(11):1860–1877, 2019.

34 Fábio K Mendes and Matthew W Hahn. Why concatenation fails near the anomaly zone. Systematic biology, 67(1):158–169, 2018.

35 Sebastien Roch, Michael Nute, and Tandy Warnow. Long-branch attraction in species tree estimation: inconsistency of partitioned likelihood and topology-based summary methods. Systematic biology, 68(2):281–297, 2019.

36 Sebastien Roch and Mike Steel. Likelihood-based tree reconstruction on a concatenation of aligned sequence data sets can be statistically inconsistent. Theoretical population biology, 100:56–62, 2015.

37 Edith M Ross and Florian Markowetz. Onconem: inferring tumor evolution from single-cell sequencing data. Genome biology, 17(1):69, 2016.

38 Erfan Sadeqi Azer, Farid Rashidi Mehrabadi, Salem Malikić, Xuan Cindy Li, Osnat Bartok, Kevin Litchfield, Ronen Levy, Yardena Samuels, Alejandro A Schäffer, E Michael Gertz, et al. Phiscs-bnb: a fast branch and bound algorithm for the perfect tumor phylogeny reconstruction problem. Bioinformatics, 36(Supplement_1):i169–i176, 2020.

39 Sohrab Salehi, Fatemeh Dorri, Kevin Chern, Farhia Kabeer, Nicole Rusk, Tyler Funnell, Marc J Williams, Daniel Lai, Mirela Andronescu, Kieran R Campbell, et al. Cancer phylogenetic tree inference at scale from 1000s of single cell genomes. Peer Community Journal, 3, 2023.

40 Gryte Satas, Simone Zaccaria, Geoffrey Mon, and Benjamin J Raphael. Scarlet: single-cell tumor phylogeny inference with copy-number constrained mutation losses. Cell systems, 10(4):323–332, 2020.

41 Susanne Schulmeister. Inconsistency of maximum parsimony revisited. Systematic biology, 53(4):521–528, 2004.

42 Michael A Steel, Michael D Hendy, and David Penny. Parsimony can be consistent! Systematic biology, 42(4):581–587, 1993.

43 David L Swofford, Peter J Waddell, John P Huelsenbeck, Peter G Foster, Paul O Lewis, and James S Rogers. Bias in phylogenetic estimation and its relevance to the choice between parsimony and likelihood methods. Systematic biology, 50(4):525–539, 2001.

44 Naoko Takezaki and Masatoshi Nei. Inconsistency of the maximum parsimony method when the rate of nucleotide substitution is constant. Journal of molecular evolution, 39(2):210–218, 1994.

45 Cuong V Than and Noah A Rosenberg. Consistency properties of species tree inference by minimizing deep coalescences. Journal of Computational Biology, 18(1):1–15, 2011.

46 Aad W Van der Vaart. Asymptotic statistics, volume 3. Cambridge university press, 2000.

47 Yong Wang and Nicholas E Navin. Advances and applications of single-cell sequencing technologies. Molecular cell, 58(4):598–609, 2015.

48 Leah L Weber and Mohammed El-Kebir. Distinguishing linear and branched evolution given single-cell dna sequencing data of tumors. Algorithms for Molecular Biology, 16(1):14, 2021.

49 Yi-Chieh Wu, Matthew D Rasmussen, Mukul S Bansal, and Manolis Kellis. Most parsimonious reconciliation in the presence of gene duplication, loss, and deep coalescence using labeled coalescent trees. Genome research, 24(3):475, 2014.

50 Yufeng Wu. Accurate and efficient cell lineage tree inference from noisy single cell data: the maximum likelihood perfect phylogeny approach. Bioinformatics, 36(3):742–750, 2020.

51 Qing Yang, Yuheng Liu, Jijian Yang, Xing Wu, Zhirui Yang, Yonghe Xia, Yunchao Zheng, Jinhong Lu, Mengdie Yao, Yiheng Du, et al. Phylosolid: Robust phylogeny reconstruction from single-cell data despite inherent error and sparsity. bioRxiv, pages 2026–02, 2026.

52 Hamim Zafar, Anthony Tzen, Nicholas Navin, Ken Chen, and Luay Nakhleh. Sifit: inferring tumor trees from single-cell sequencing data under finite-sites models. Genome biology, 18(1):178, 2017.

